# Transcriptional landscape of the dorsal raphe serotonin neurons rendering stress resiliency

**DOI:** 10.1101/2024.03.21.586199

**Authors:** Chihiro Andoh, Suzuka Otani, Takuma Noguchi, Masako Hagiwara, Naoya Nishitani, Hiroyuki Kawai, Yuto Fukui, Masashi Koda, Hinako Morishita, Kento Nomura, Moeka Oki, Harune Hori, Hisashi Shirakawa, Shuji Kaneko, Kazuki Nagayasu

## Abstract

Major depressive disorder (MDD) is a serious and large social problem, yet the pathophysiology of MDD and the action mechanism of antidepressants are still poorly understood. A number of studies have reported that activation and inactivation of serotonin neurons in the dorsal raphe nucleus (DRN) cause antidepressant-like effects and depressive-like behaviors, respectively. Also, their physiological neural activities are increased when mice were chronically administered an SSRI and decreased in mice exposed to chronic social defeat stress (CSDS), a mouse model of depression. However, the molecular mechanism underlying these neural activity changes in DRN serotonin neurons remains unclear. In this study, we performed a DRN serotonin neuron-specific comprehensive gene expression analysis by using Translating Ribosome Affinity Purification (TRAP) technology in both chronic SSRI-treated mice as a model of antidepressant treatment and CSDS mice as a model of depression. It revealed that many gene expression changes were the opposite between SSRI-treated mice and CSDS-susceptible mice. Among these, we identified S100a10 as a prodepressive gene in DRN serotonin neurons, and we found that Interleukin-4 (IL-4) – Signal Transducer and Activator of Transcription 6 (STAT6) pathway and 5-HT_1B_ receptor were the upstream and downstream molecules of S100a10, respectively. Our findings provide insights into molecular mechanisms underlying the action of antidepressants and stress resiliency.

---

Major depressive disorder (MDD) is a severe and sizeable social problem, affecting more than 300 million people worldwide ^1,2^. Environmental stressors are one of the most crucial risk factors for MDD ^3^. While adequate stress stimulus promotes adaptation to physical or psychological threats ^4^, prolonged exposure to excessive stress leads to brain function deterioration and, ultimately MDD ^5^. However, the neural substrates essential for successful adaptation to stress are not yet fully understood. Numerous reports have demonstrated the critical role of the serotonin system originating from the dorsal raphe nucleus (DRN) in stress resiliency as well as stress coping ^6^. Anatomically, the DRN is the largest serotonergic nuclei reciprocally interconnected with critical brain regions underlying increase, decrease, and maintenance of stress resiliency, including the medial prefrontal cortex, amygdala, lateral habenula, and ventral tegmental area ^7–11^, suggesting the critical hub role of DRN orchestrating brain regions rendering stress resiliency. Moreover, we and others have demonstrated that the activity of DRN serotonin neurons is decreased by chronic stress and is increased by chronic treatment with antidepressants, which renders stress resiliency ^12–17^. Mechanistically, we and others have reported that repeated optogenetic activation of the DRN serotonin neurons induces stress resiliency, while their chronic inhibition leads to loss of stress resiliency ^18,19^. Although these results strongly suggest the deterministic role of the activity of DRN serotonin neurons in stress resiliency, the mechanisms underlying the chronic activity changes of DRN serotonin neurons are yet to be unveiled.

In this study, we investigated how the transcriptomic landscape of DRN serotonin neurons changes after spontaneously acquired stress resiliency (resilient mice after chronic social defeat stress) and drug-induced stress resiliency (chronic antidepressant). We identified the downregulation of S100a10 in both stress resiliency models and its causal relationships with antidepressant-like effects. Unexpectedly, upstream analysis of altered transcriptomic landscape revealed a critical role of IL-4 and its downstream signal transducer STAT6, well-known type 2 helper T-cell (Th2) cytokine signaling ^20–23^, in both stress resiliency models. Chronic treatment with SSRIs induced a transient increase in IL-4 level in the DRN and phosphorylated STAT6 in the DRN serotonin neurons. A transient application of IL-4 to the DRN induced an antidepressant-like effect, while attenuation of IL-4 signaling by dominant-negative STAT6 in DRN serotonin neurons resulted in the opposite effect. Our study unveiled the dynamic changes of transcriptomic landscapes in the DRN serotonin neurons and how these changes render stress resiliency under natural stress adaptation and antidepressant treatment, as well as a critical role of IL-4-STAT6 signaling in the acquisition of stress resiliency.

## Transcriptome landscape of the DRN serotonin neurons

To evaluate the transcriptome of the DRN serotonin neurons, we performed the translating ribosome affinity purification (TRAP) through selective expression of GFP-tagged ribosomal subunit (EGFP-Rpl10A) ^24–27^. We confirmed selective expression of EGFP-Rpl10A in the DRN serotonin neurons four weeks after injection of the adeno-associated viral vectors (AAVs) bearing EGFP-Rpl10A under Tph2 promoter^19,28,29^ (90.2 ± 4.26% coverage, 85.8 ± 2.59% specificity; Fig.1a, Extended Data Fig.1a). Whole-tissue (input) and immunoprecipitated RNA (TRAP) was extracted from the DRN (Fig. 1b; n = 3 groups of three mice), yielding sufficient amount of intact RNA (Extended Data Fig.3a, b). Microarray analysis revealed the enriched expression of serotonin neuron markers, including *Tph2*, *Ddc*, and *Slc6a4*, in the TRAP samples, while the expression of glial cell markers, including *Gfap*, *Mbp*, and *Aif1* (Iba1), were depleted (Fig. 1c, Extended Data Fig. 1b, c, Extended Data Table 1), similar to previous studies in other types of neurons^30,31^.

**Figure 1.**
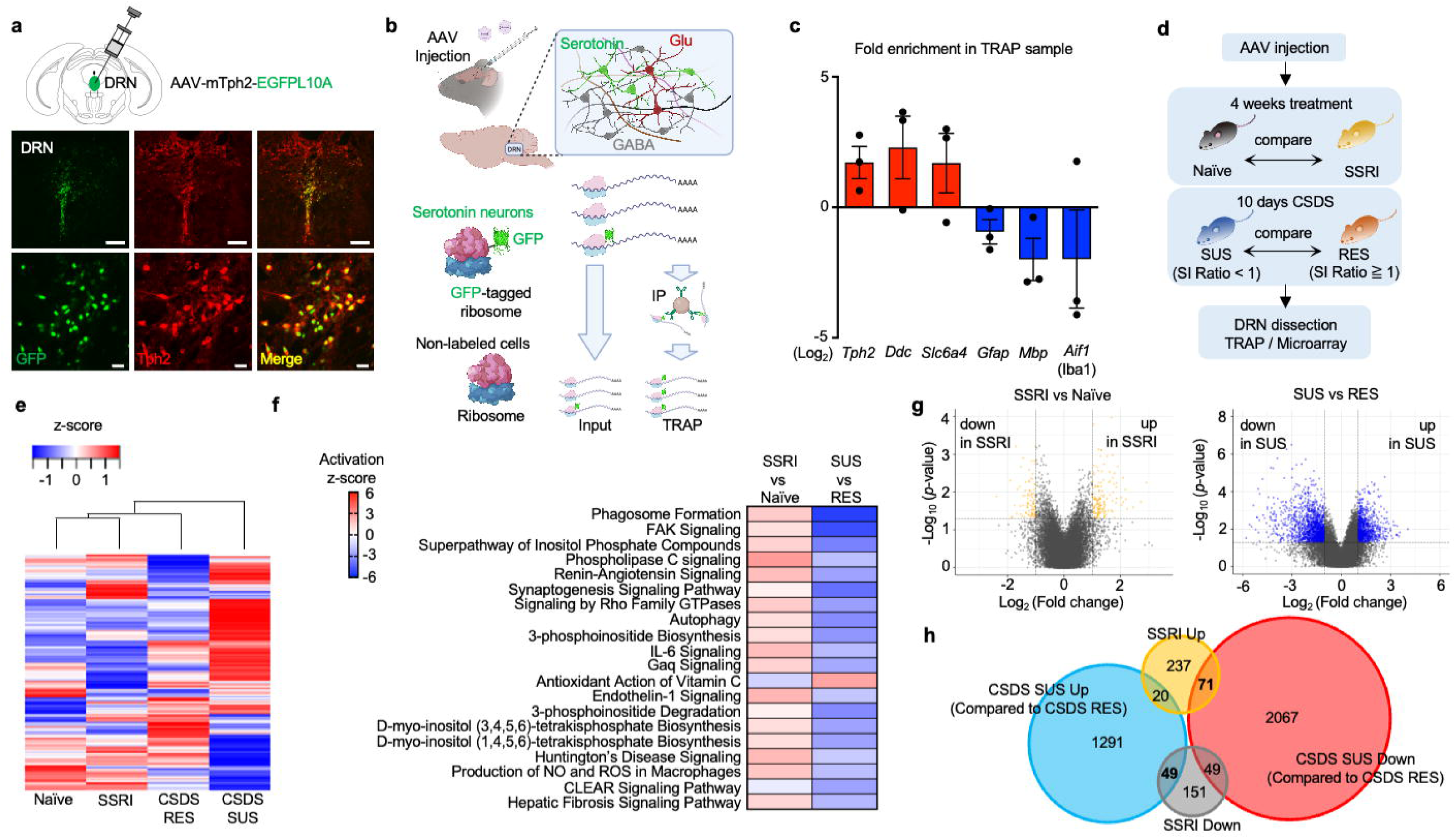
Serotonin neuron-specific translatome in SSRI treated mice and CSDS mice. (a) Adeno Associated Virus vector construction (top) and immunofluorescence image of viral expression in DRN (bottom). Scale bars = 200 μm (top panel), 20 μm (bottom panel). (b) Schematic image of TRAP technology. (c) Fold enrichment of serotonin neuron markers and glial cell markers in TRAP sample compared to Input sample. n = 3 and 3 mice were pooled for each sample. (d) Schematic overview and time course of the experiments. (e) Hierarchical clustering heatmap of gene expression. Z-score was calculated from their expression values (log_2_). Each group contains 3 sample and 2 to 3 mice were pooled for each sample. (f) IPA comparison analysis of canonical pathways. A top 20 pathways are displayed in this graph. (g) Gene expression analysis between naïve mice and SSRI treated mice (left) and between CSDS resilient mice and susceptible mice (right). Colored dots indicate genes with more than 2-fold expression changes and p-value with less than 0.05. (h) Venn diagram of differential expression genes (DEGs). DEGs were defined as genes whose expression changes were 2-fold or more.

To investigate the molecular mechanism in the DRN serotonin neurons underlying the acquisition of stress resiliency, we performed TRAP analysis of the DRN serotonin neurons in chronic vehicle-treated mice (naïve), chronic SSRI-treated mice (citalopram in drinking water (∼24 mg/kg/day) for 28 days), and CSDS mice. Four weeks of citalopram administration in drinking water (∼24 mg/kg/day) increased active-coping behaviors in the tail-suspension test (TST) without affecting the locomotor activity in the open-field test (OFT) in an independent cohort of the animals used in the TRAP analysis to avoid any confounding effect of the behavioral tests. (Extended Data Fig. 2a, b). CSDS mice were divided into two groups based on the social interaction (SI) ratio (CSDS susceptible (SI ratio < 1) and CSDS resilient (SI ratio ≥ 1)), an index of stress resiliency, according to the previously reported criteria (Golden et al., 2011, Extended Data Fig. 2c-f). Then, we compared the gene expression changes between naïve and chronic SSRI-treated mice with those between CSDS susceptible and resilient mice to identify common gene expression changes during the acquisition of stress resiliency (Fig.1d, Extended Data Fig. 3a, b, Extended Data Table 2). The gene expression of the same treatment group generally showed a strong correlation (Extended Data Fig.3c), indicating high repeatability. We noted that one sample in the CSDS resilient group (CSDS RES-1) showed gene expression similar to that of the CSDS susceptible group. Interestingly, the mean SI ratio in the CSDS RES-1 group was the lowest among the CSDS resilient groups, at 1.53. Moreover, previous reports indicate that the SI ratio of naïve mice is around 1.5, although the commonly used threshold between CSDS resilient and susceptible groups is an SI ratio of 1.0^32,33^. Although these data may indicate that mice showing an SI ratio of around 1.5 should be considered as a CSDS susceptible group, we followed the conventional criteria in this study for consistency with previous reports. The hierarchical clustering analysis of overall gene expressions revealed that CSDS-susceptible mice showed the most different gene expressions from the other groups (Fig. 1e). Pathway analysis using QIAGEN ingenuity pathway analysis (IPA) revealed that the pathway activated in the SSRI group was inhibited in the CSDS susceptible group (Fig. 1f), suggesting that acquisition of stress resiliency caused similar transcriptome changes in the DRN serotonin neurons. Nevertheless, the number of differentially expressed genes (DEGs) between CSDS susceptible and resilient groups was far greater than that between SSRI and naïve groups (Fig. 1g). We identified 49 genes whose expressions were increased in CSDS susceptible groups and decreased by chronic SSRI treatment and 71 genes whose expressions were increased by SSRI and decreased by CSDS (Fig. 1h), as a possible determinant of stress resiliency.

## Decreased expression of S100a10 induces an antidepressant-like effect through downregulation of 5-HT_1B_ receptors in the DRN serotonin neurons

We analyzed the enrichment of the identified genes in serotonin neurons by comparing the expression levels in the input and TRAP samples (Fig. 2a, b). The most enriched gene was *Gchfr* (7.26 fold enrichment), coding GTP cyclohydrolase I feedback regulator, which was involved in the synthesis of tetrahydrobiopterin ^34^; however, mere knockdown of *Gchfr* does not affect the activity of GTP cyclohydrolase and the level of tetrahydrobiopterin^35^ and the expression changes of *Gchfr* after SSRI treatment (0.45 fold change) and CSDS (2.31 fold change) was marginal (Extended Data Table 2). We then focused on the second most enriched gene in serotonin neurons, *S100a10* (5.98 fold enrichment), whose expression decreased in SSRI-treated mice (0.32 fold change) and increased in CSDS-susceptible mice (2.14 fold change). S100a10 is a member of the S100 family protein ^36^ and is widely expressed in humans and mice (Marenholz et al., 2004; Saris et al., 1987). Although it is reported that S100a10 is expressed not only in neurons but also in glial cells and ependymal cells ^38^, immunohistochemical analysis revealed that S100a10 was expressed almost selectively in serotonin neurons in the DRN (95.3 ± 0.92% specificity; Extended Data Fig. 4). Although S100a10 in other brain regions such as the nucleus accumbens (NAc) and hippocampus is involved in depressive-like behavior and antidepressant effects in mice and humans^39–42^, the role of S100a10 in DRN serotonin neurons is still unknown. Immunohistochemical analysis in a different cohort of mice revealed that the protein expression of S100a10 in DRN decreased after four weeks of SSRI treatment and increased in the CSDS susceptible mice compared to resilient mice (Fig. 2c, d, Extended Data Fig.5). Western blotting analysis of whole DRN tissue showed the decreased tendency of S100a10 expression in chronic SSRI-treated mice (Extended Data Fig. 6). Knockdown of S100a10 in the DRN serotonin neurons (Fig. 2e-g) increased active coping behaviors in the TST without affecting locomotor activity in the OFT and spent time in the open arm in the elevated plus maze test (EPM) (Fig. h-j).

**Figure 2.**
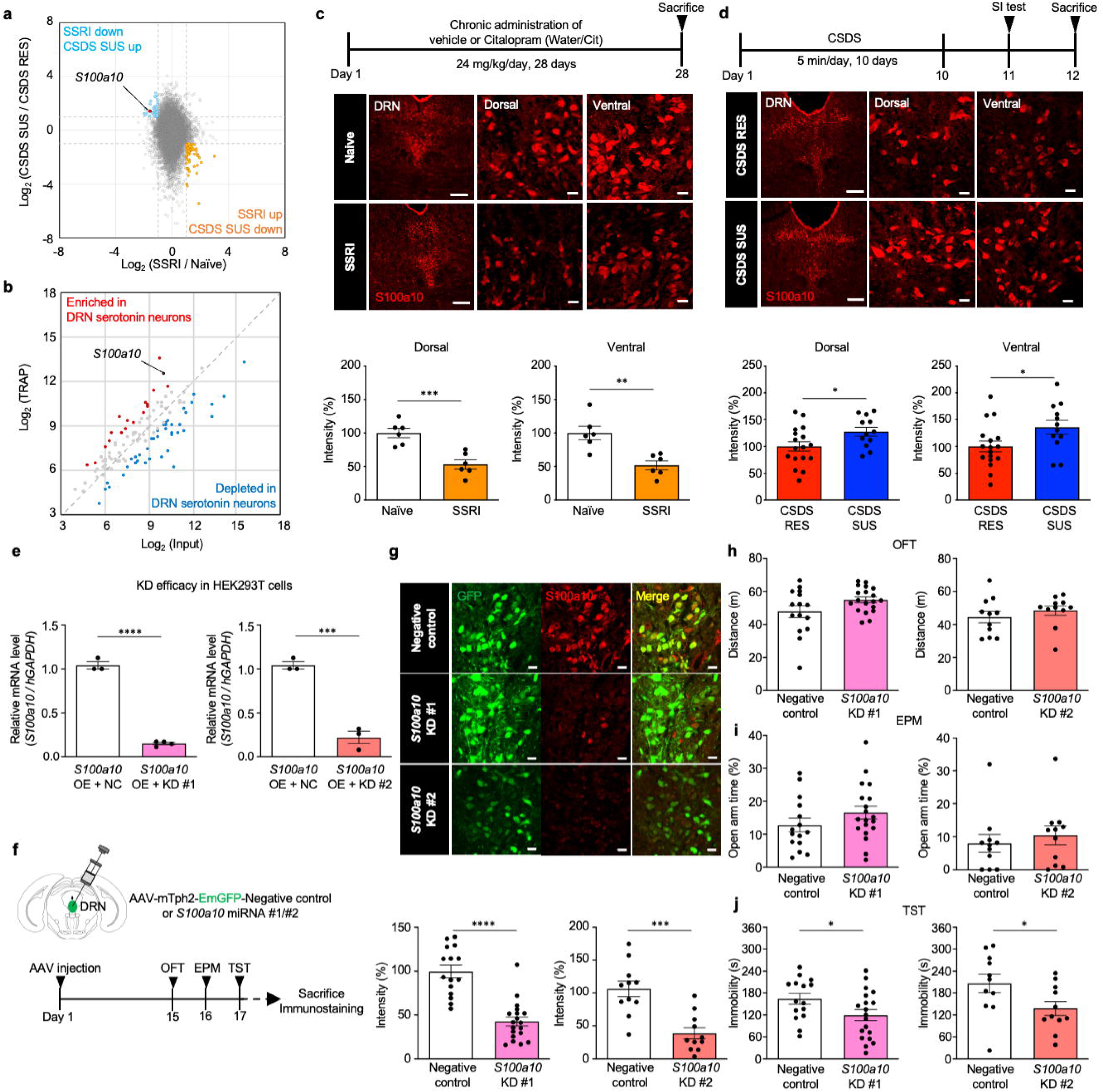
DRN serotonin neuron specific knockdown of S100a10 induces an antidepressant-like effect. (a) The graph represents the expression changes of the SSRI treated mice compared to the naïve mice on the horizontal axis and the expression changes of the CSDS susceptible mice compared to the CSDS resilient mice on the vertical axis. Light blue dots were the genes whose expression changes were significantly increased in CSDS susceptible mice and decreased in SSRI treated mice, whereas the orange dots vice versa. (b) The serotonin neuron enrichment of significant genes in (a). Red dots were the genes whose expressions were enriched 2-fold or more in the TRAP sample and blue dots were the genes whose expressions were depleted 2-fold or more in the TRAP sample compared to Input. (c) Time course of SSRI treatment (top), immunofluorescence image (middle) and quantification of S100a10 (bottom) in DRN of naïve mice and SSRI treated mice. Scale bar, 200 μm (left panel), 20 μm (middle and right panels). n = 6. \*\**P* < 0.01 and \*\*\**P* < 0.001, two-tailed t-test. (d) Time course of CSDS (top), immunofluorescence image (middle) and quantification of S100a10 (bottom) intensity in DRN of CSDS resilient and susceptible mice. Scale bar, 200 μm (left panel), 20 μm (middle and right panels). n = 12-17. \**P* < 0.05, two-tailed t-test. (e) S100a10 mRNA expression in Lenti-X 293T cells treated with the negative-control plasmid or the S100a10 knockdown plasmid. The expression was normalized by hGAPDH. n = 4. \*\*\*\**P* < 0.0001, two-tailed t-test. (f) Schematic representation of AAV injection and time course of experiments. (g) Immunofluorescence image (top) and quantification of S100a10 intensity (bottom) in DRN of control and knockdown mice. See Materials & Methods for the sequence used for each knockdown. Scale bar, 20 μm. n = 15-19 (for KD #1 experiments), 11 (for KD #2 experiments). \*\*\**P* < 0.0001, \*\*\*\**P* < 0.0001, two-tailed t-test. (h) Total distance in Open Field Test (OFT), (i) time spent in open arm in Elevated Plus Maze test (EPM) and (j) total immobility time in Tail Suspension Test (TST). n = 15-19 (for KD #1 experiments), 11 (for KD #2 experiments). \**P* < 0.05, two-tailed t-test. All data presented as means ± s.e.m.

Lines of reports have demonstrated that S100a10 interacts with many ion channels, GPCRs, and enzymes and modulates their functions by mainly changing their localization ^43^. This interaction can be divided into two types depending on the presence or absence of annexin A2 (AnxA2) dependence ^44^. Since our TRAP data and a previous report^45^ suggested that *Anxa2* is weakly expressed in the serotonin neurons (Extended Data Table 1), we focused on the molecules with which S100a10 interacts even in the absence of AnxA2. The candidate molecules were three serotonin receptor subtypes (5-HT_1B_, 5-HT_1D_, and 5-HT_4_ receptors), CC-chemokine receptor 10 (CCR10), acid-sensing ion channel 1a (ASIC1a), and TWIK-related acid-sensitive K^+^ channel 1 (TASK1) ^46–50^. Among these, we focused on 5-HT_1B_ receptors, most abundantly expressed in the DRN serotonin neurons (Extended Data Table 1). Previous reports indicate that depletion of S100a10 abolishes the function of 5-HT_1B_ receptors ^17,46^. Interestingly, 5-HT_1B_ receptors in the serotonin neurons inhibit serotonergic neurotransmission as an inhibitory autoreceptor^51^. Then, we evaluated the 5-HT_1B_ receptor expression levels in S100a10 knockdown mice. Immunohistochemical analysis revealed that 5-HT_1B_ receptor expression levels were significantly decreased by S100a10 knockdown in the DRN serotonin neurons (Fig. 3a). Moreover, serotonin neuron-selective knockdown of 5-HT_1B_ receptors in the DRN (Fig. 3b-d) significantly increased active coping behaviors in the TST without affecting locomotor activity in the OFT and spent time in the open arm in the EPM (Fig. 3e-g). These results indicate that decreased expression of S100a10 under the acquisition of stress resiliency is causally related to the induction of antidepressant-like effects through the downregulation of inhibitory 5-HT_1B_ autoreceptors in the DRN serotonin neurons.

**Figure 3.**
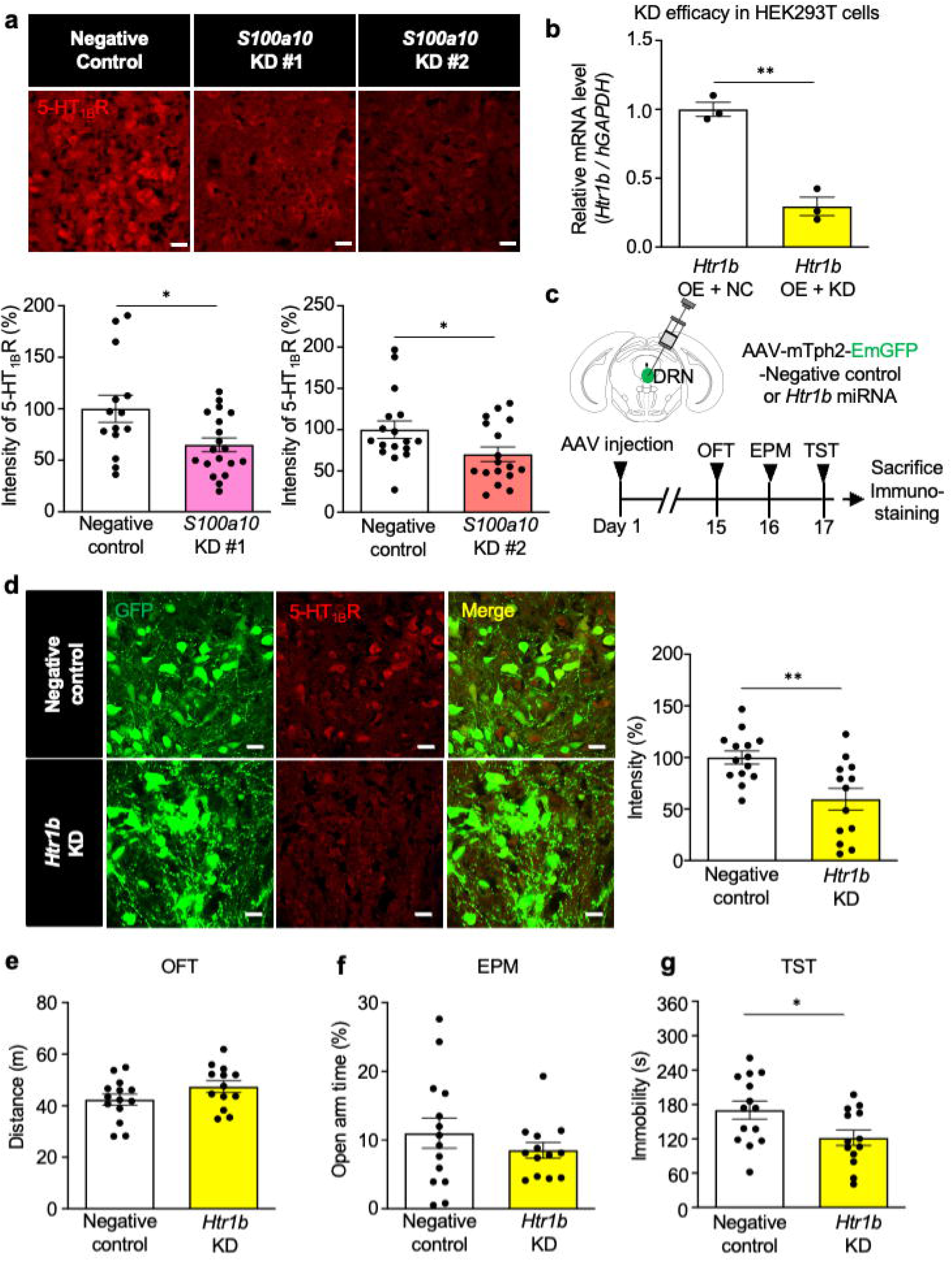
DRN serotonin neuron specific knockdown of 5-HT_1B_R also shows an antidepressant-like effect. (a) Immunofluorescence image (top) and quantification of 5-HT_1B_R (bottom) in DRN of control mice and S100a10 knockdown mice. Images were taken in areas of the DRN where AAV vector was well expressed (mainly ventral part of the DRN). Scale bar, 20 μm. n = 14-19 (for KD #1 experiments), 17 (for KD #2 experiments). \**P* < 0.05, two-tailed t-test. (b) Htr1b mRNA expression in HEK 293T cells treated with the negative-control plasmid or the Htr1b knockdown plasmid. The expression was normalized by hGAPDH. n = 4. **P < 0.01, two-tailed t-test. (c) Schematic representation of AAV injection and time course of experiments. (d) Immunofluorescence image (left) and quantification of 5-HT_1B_R intensity (right) in DRN of control and knockdown mice. Scale bar, 20 μm. n = 13-14. **P < 0.01, two-tailed t-test. (e) Total distance in OFT, (f) time spent in open arm in EPM, (g) total immobility time in TST. n = 13-14. *P < 0.05, two-tailed t-test. All data presented as means ± s.e.m.

## Activation of the IL-4-STAT6 pathway in the DRN serotonin neurons under the acquisition of stress resiliency

We sought the mechanism underlying the decreased expression of S100a10 through pathway analysis of transcriptome changes under the acquisition of stress resiliency. We identified seven molecules upstream to S100a10 through ingenuity pathway analysis (IPA) of the transcriptome changes (Fig. 4a). Unexpectedly, we found that the IL-4 and its downstream signal transducer STAT6 pathway (Fig. 4b), well-known type 2 helper T-cell (Th2) cytokine signaling in immune system ^20–23^, showed opposite and robust activation states in SSRI-treated mice and CSDS-susceptible mice.

**Figure 4.**
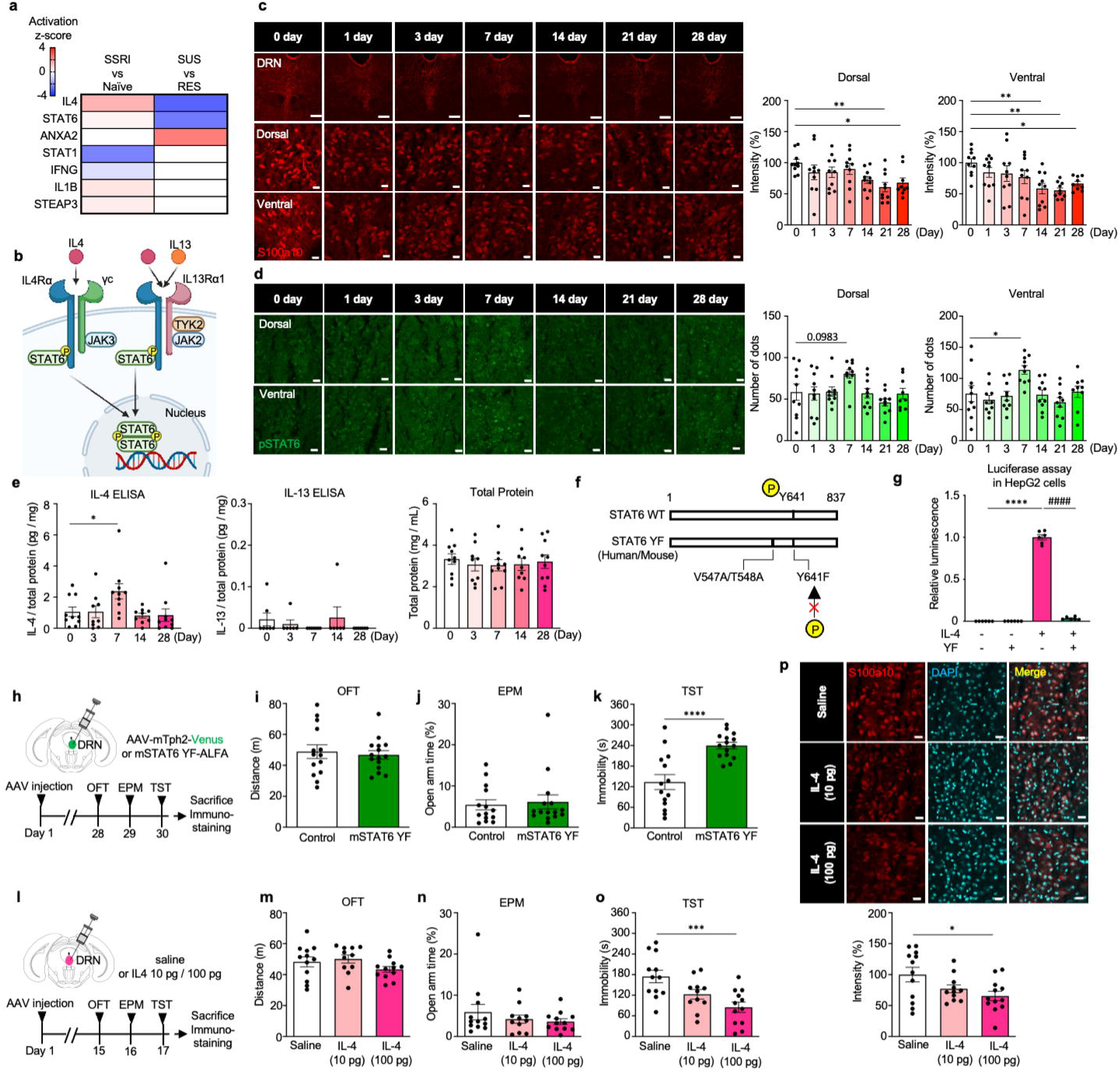
Activation of IL-4-STAT6 pathway in DRN serotonin neurons induces an antidepressant-like effect via downregulation of S100a10. (a) IPA comparison analysis of upstream pathways. (b) A schematic illustration of the IL4-STAT6 signaling pathway. (c) Immunofluorescence image (left) and quantification of S100a10 (right) in DRN of SSRI-treated mice at each time points. Scale bar, 200 μm (top pannel), 20 μm (middle and bottom panels). n = 9-10. **P < 0.01, compared to 0 day, one-way ANOVA, Dunnett’s *post hoc* test. (d) Immunofluorescence image (left) and quantification of pSTAT6 (phosphor Y641) (right) in DRN of SSRI-treated mice at each time point. Scale bar, 20 μm. n = 9-10. *P < 0.01, compared to 0 day, one-way ANOVA, Dunnett’s *post hoc* test. (e) ELISA measurements for IL-4 (left) and IL-13 (middle) levels of DRN at each time point after SSRI treatment. Total protein levels of DRN samples measured by BCA assay (right). n = 9-10. Data presented as means ± s.e.m. n = 9-10. *P < 0.01, compared to 0 day, one-way ANOVA, Dunnett’s *post hoc* test. (f) An illustration of the STAT6 mutant used in the following experiments. (g) Luciferase activity of 48 h posttransfection of hSTAT6 YF or pcDNA in either unstimulated HepG2 cells or cells that had been treated with IL-4 6 h prior to harvest. We conducted two independent assays (n = 3 each) and the luciferase activity was normalized by each batch of the IL-4 (+) YF (-) group. ****P < 0.0001, compared to IL-4 (-) YF (-), ####P < 0.0001, compared to IL-4 (+) YF (-), respectively by One-way ANOVA, Tukeys *post hoc* test. (h) Schematic representation of AAV injection and time course of experiments. (i) Total distance in OFT, (j) time spent in open arm in EPM, (k) total immobility time in TST. n = 14-15. ****P < 0.0001, two-tailed t-test. (l) Schematic representation of drug injection and time course of experiments. (m) Total distance in OFT, (n) time spent in open arm in EPM, (o) total immobility time in TST. n = 11-12. ***P < 0.001, compared to saline, one-way ANOVA, Dunnett’s *post hoc* test. (p) Immunofluorescence image (top) and quantification of S100a10 (bottom) in DRN of IL4-injected mice at each time points. Scale bar, 20 μm. n = 11-12. *P < 0.05, compared to saline, one-way ANOVA, Dunnett’s *post hoc* test. All data presented as means ± s.e.m.

Immunohistochemical analysis revealed the transient increase of phosphorylated STAT6 (pSTAT6) in the DRN serotonin neurons after 7 days of SSRI treatment, followed by the sustained decrease in S100a10 expression after SSRI treatment for 14 days and more (Fig. 4c, d, Extended Data Fig.7). Western blotting of the whole DRN tissue showed that pSTAT6/STAT6 ratio tended to increase after 7 days of SSRI treatment (Extended Data Fig. 8). Consistent with the result that pSTAT6 level reversed to the baseline when S100a10 expression decreased in chronic SSRI-treated mice, we found no significant difference in the pSTAT6 level between CSDS-susceptible and CSDS-resilient groups (Extended Data Fig. 9).

STAT6 is primarily stimulated by IL-4 and IL-13 ^52^, and we found that both IL-4Rα and IL-13Rα1 were abundantly expressed in the TRAP samples compared to the expression of common gamma chain (Extended Data Table 1). This result was consistent with the previous report that the IL-4 type 2 receptor (IL-4Rα and IL-13Rα1 complex) is mainly expressed in non-hematopoietic cells, while IL-4 type 1 receptor (IL-4Rα and common gamma chain complex) is expressed primarily in hematopoietic cells ^53^. The levels of IL-4 and IL-13 in the whole DRN tissue were analyzed by ELISA to determine whether and when these cytokines increased during chronic SSRI treatment. We found a transient increase of IL-4 on day 7 of chronic SSRI treatment, the same time point of STAT6 activation. In contrast, IL-13 was almost undetectable in all time points (Fig. 4e). These results indicated that IL-4-STAT6 pathway, not IL-13, may play a vital role in an antidepressant-like effect. To further elucidate the mechanism, we first attempted to inhibit this pathway by using a mutant of STAT6 with the replacement of Tyr-641 by a phenylalanine residue (STAT6 YF, Fig.4f). This mutant is inactive because it is not phosphorylated. Luciferase assays from HepG2 cells transfected with the mutant expression plasmids showed that the overexpression of the mutant suppressed the endogenous STAT6 activity in response to IL-4 (Fig.4g). Therefore, we produced the AAV which serotonin neuron specifically expressed this mutant STAT6 and conducted behavioral analysis (Fig.4h). As a result, mice expressed mutant STAT6 showed significantly increased immobility time in TST, while there was no difference in locomotor and anxiety (Fig.4i-k). These results indicated that STAT6 inhibition led to a pro-depressive like effect. Furthermore, we investigated whether IL-4 microinjection in DRN is sufficient to cause an antidepressant-like effect. While locomotor behavior and anxiety were not different between all groups, the immobility time in TST was decreased in a concentration-dependent manner, and mice with 100 pg of IL-4 injection showed significantly short immobility time compared to control mice (Fig.4l-o). Also, the expression of S100a10 was significantly decreased in 100 pg IL-4 injection mice (Fig.4p), with increased pSTAT6 expression immediately after IL-4 injection (Extended Data Fig.10). Taken together, these results indicated that IL-4–STAT6 pathway in DRN serotonin neurons plays a crucial role in an antidepressant-like effect via regulation of S100a10 expression.

## Discussion

In this study, we investigated the molecular mechanisms underlying the action of antidepressants and stress resiliency in the DRN serotonin neurons by comprehensive gene expression analysis of chronic SSRI-treated model mice and CSDS model mice. An advantage of the CSDS model is that it can separate susceptible and resilient mice under identical stress conditions^54^ and is thus ideal for elucidating the mechanisms of stress resiliency more accurately than previous studies that solely compared gene expression changes under stressed and unstressed conditions by using LPS-induced depression model^55^ or forced swim stress model^56^. In detail, Lesiak *et al.* ^56^ conducted a comprehensive gene expression analysis of the DRN serotonin neurons under the condition of two days of forced swim stress using ePet-Cre^tg/−^/RiboTag^tg/−^ mice. They identified that Fkbp5 was enriched in serotonergic neurons and upregulated by stress, and they also reported that inhibition of Fkbp5 in the DRN blocked stress-induced reduction in sucrose consumption. Our TRAP data also showed the significant upregulation of Fkbp5 in CSDS susceptible mice compared to naïve mice, but we found expressions of the gene were also upregulated in SSRI-treated mice and CSDS resilient mice (Extended Data table 1), suggesting that this gene is not fully illustrate a mechanism of stress resiliency. Therefore, in this study, we compared the gene expression of CSDS susceptible and resilient mice like the previous reports, which defined a part of the molecular mechanism of stress resiliency in the nucleus accumbens or hippocampus using CSDS mice ^57,58^. Furthermore, we directly compared gene expression changes between the CSDS model and the antidepressant treatment model and identified the candidate pro-depressive gene *S100a10*.

Apparently contradicting our result, it has been reported that S100a10 knockout mice showed depressive-like symptoms ^46^ and S100a10 protein and mRNA in the cortex and hippocampus are known to increase after SSRI administration and decrease in both a mouse model of depression and depressive patients ^40,59–62^. On the other hand, S100a10 was significantly upregulated by chronic restraint stress in the lateral habenula ^63^, the same as our result of DRN serotonin neurons. These apparent discrepancies may be due to the different mechanisms by which S100a10 regulates neural activity in these brain regions through its interactions with molecules such as SMARCA3 and mGluR5 ^41,64,65^.

In this study, we have presented the possibility that S100a10 in DRN serotonin neurons affected the antidepressant effects by decreasing the amount of 5-HT_1B_R on the membrane. 5-HT_1B_R act as autoreceptors primarily at the axon terminals^66^ but have also been reported to be functionally expressed in the cell bodies of DRN serotonin neurons^67,68^. Indeed, a previous report has shown that 5-HT_1B_R activation in the DRN decreases the release of serotonin^69^, and serotonin neuron-specific knockout of 5-HT_1B_R caused an antidepressant-like effect without drug administration^70^, which supports our current results. Surprisingly, we found the decreased expression of 5-HT_1B_R in the DRN of S100a10 knockdown mice, even though S100a10 was predicted to be involved only in membrane trafficking of 5-HT_1B_R and not to affect its expression level. This decrease may be due to the degradation of internalized GPCR in lysosomes through ubiquitination or other pathways^71–73^. Collectively, our study uncovered the role of S100a10 in DRN serotonin neurons, one of the most critical components of the pathophysiology of MDD.

Several transcription factors such as activator protein-1 (AP-1) complex formed by c-Fos and c-Jun, and GRHL2, a grainyhead-like transcription factor family member, have been reported as upstream regulators of S100a10^74,75^. In this study, we identified the transient activation of IL4-STAT6 pathway regulates the expression of S100a10 in the DRN serotonin neurons. IL-4 is the well-known type 2 helper T-cell (Th2) cytokine, although it is also produced by macrophages, mast cells, central neurons and M2-polarized microglia ^53,76,77^. Numerous studies have revealed the relationship between cytokines and the pathophysiology of depression^78,79^. However, most of these studies were focused on Th1 cytokines like TNFα and IFN-γ, which have been reported to increase in MDD patients^80–82^ and mice model of depression^83^. For example, the release of TNFα was stimulated through the activation of NF-κB signaling^84^, and TNFα in the hippocampus decreased the hippocampal norepinephrine level and decreased brain-derived neurotrophic factor (BDNF), which leads to the development of depression^85^. Also, TNFα has been reported to disrupt the glucocorticoid receptor signaling and contribute to altered glucocorticoid-mediated feedback regulation^86^. Compared to these abundant reports on Th1 cytokines, the precise role of Th2 cytokines is still unclear, although some reports implicated that serum IL-4 levels were decreased in MDD patients ^87^ and recovered by SSRI treatment^88^. Considering that an imbalance between Th1/Th2 cytokines is thought to be a potential risk factor for depression^89^, it is suggested that induction of IL-4 in DRN during SSRI treatment found in this study may restore the Th1/Th2 balance and thus result in antidepressant effects. Indeed, a meta-analysis indicated that anti-inflammatory drugs such as corticosteroids and NSAIDs have a therapeutic impact on depression^90^, although another report found the effect was negligible^91^. These drugs have been reported to inhibit the production of Th1 cytokines such as TNFα and IFNγ^92^. On the other hand, these drugs have been reported not only to decrease the production of Th2 cytokines under allergic conditions^93^ but also to increase Th2 cytokines by acting on splenic adherent cells (SpAC) to promote differentiation of uncommitted T cells into Th2 cells via suppression of IL-12^94^. The balance of which cells the drugs work effectively on may alter its impact on depression. Also, some preclinical reports suggested the protective role of IL-4 in the pathophysiology of depression. For example, IL-4 deficient mice showed an increase of depressive-like behavior ^95^, while mice resilient to chronic mild stress showed higher hippocampal IL-4 and the microinjection of IL-4 in this region strengthened the resiliency to stress ^58^. Furthermore, intranasal administration of IL-4 ameliorated depressive-like behavior through the modulation of neuroinflammation and oxidative stress ^96^, all of which are consistent with our findings in this report. One of the remaining questions is that which cells produce IL-4 in the DRN. Given that blood-brain barrier (BBB) disruption is considered to be occurred in MDD patients and CSDS model mice ^97,98^ and some peripheral immune cells including T cells can enter the brain parenchyma even under the physiological condition^99,100^, IL-4 produced by peripheral Th2 cells possibly contribute to the anti-depressant like effect in DRN, although we could not exclude the possibility of IL-4 production by CNS cells such as M2-polarized microglia or serotonin neurons themselves. Further immunohistological analysis will be needed to elucidate the precise mechanism.

In the experiment using bone marrow-derived macrophages, DNA binding of STAT6 induced by IL-4 stimulation has been reported to be immediate and transient, with a steep increase in binding at 1 hour after stimulation followed by a decrease to baseline after 24 hours^101^. Nevertheless, gene expression changes (both upregulation and downregulation) induced by STAT6 occur for a delayed and prolonged period in bone marrow-derived macrophages and endothelial cells^102,103^. In this study, we found SSRI treatment induced transient activation of IL-4-STAT6 signaling in the DRN and a delayed and sustained S100a10 expression reduction, leading to an antidepressant-like effect. This may partly explain why the therapeutic effect of SSRIs is exerted only after a few weeks^104^ and why stress resiliency, once formed, lasts for more than a month^105,106^. A possible mechanism to bridge this temporal gap is epigenetics, stable and long-lasting gene expression variations induced by non-DNA-encoded mechanisms such as DNA methylation, histone modification, and RNA modification ^107,108^. It has been reported that STAT6 modulates both permissive and repressive histone modifications like H3K27ac, H3K9ac, and H3K27me3 ^101,109^. Considering that numerous studies have suggested a relationship between these histone modifications and the pathophysiology of depression^110,111^, further investigation of these histone modifications in the DRN serotonin neurons in a cell-type selective manner will provide insights into the complete picture of how antidepressants induce chronic transcriptomic changes through transient activation of IL-4-STAT6 signaling by using chromatin profiling methods applicable for a small number of cells such as CUT&Tag^112^.

In conclusion, we revealed the transcriptomic landscape of DRN serotonin neurons under the condition of antidepressant treatment and stress exposure. Among these, we identified the IL-4-STAT6-S100a10-5-HT_1B_R pathway as a crucial mediator of antidepressant-like effects, providing insights into molecular mechanisms underlying the action of antidepressants and stress resiliency. Further investigation of more detailed mechanisms will lead to a better understanding of the pathogenesis of depression and the development of rapidly-acting and safer antidepressants.

## Methods

### Animals

All animal care and experimental procedures were conducted in accordance with the ethical guidelines of the Kyoto University animal experimentation committee (approval code: 13-41-2, 19-41-1,2,3). The adult male C57BL/6J mice (6–10 weeks old, Nihon SLC, Shizuoka, Japan or maintained in our laboratory) were housed in groups unless otherwise noted. They were kept at a constant ambient temperature (22 ± 2 °C) and humidity (55 ± 10 %) under a 12 h light/dark cycle with free access to food and water. All behavioral studies were conducted in light phase.

### Drugs

For chronic antidepressant treatment, citalopram hydrobromide (a selective serotonin reuptake inhibitor; Struchem, Wujiang, China) was dissolved in saline (20 mg/mL) as a stock and stored at −20°C until use. The stock was diluted 100-fold with tap water (equal to 0.2 mg/mL) and administrated for 28 days. Water consumption was approximately 3– 4 mL/day/mouse, resulting in average dose at 24 mg/kg/day. The drug containing drinking water was shielded from light and changed every 3–5 day. When drug administration was performed on virus-injected mice (that is, mice used for TRAP or CUT&Tag), administration began at least one week after stereotaxic surgery. For the time-dependent experiment (correspond to Figure 5), the day of completion of drinking was aligned among the groups.

### Vector Construction

For construction of pAAV-mTph2-EGFPL10A-WPRE, the EGFPL10A fragment was amplified by PCR from pAAV-FLEX-EGFPL10A (addgene #98747), and ligated to AAV backbone obtained from pAAV-mTph2-GCaMP6s-WPRE ^29^.

For construction of pAAV-CMV-EmGFP-miRNA-negative control, negative control sequences (see Table below) were designed from pcDNA6.2-GW/EmGFP-miR-neg Control (Invitrogen, Carlsbad, CA, USA). For construction of pAAV-CMV-EmGFP-miRNA-*S100a10* or *Htr1b*, miRNA sequences (see Table below) targeting *S100a10* (NM_009112.2) or *Htr1b* (NM_010482.2) were designed using BLOCK-iT™ RNAi Designer (https://rnaidesigner.thermofisher.com/rnaiexpress/). Oligodeoxynucleotide primers were purchased from Sigma-Aldrich Japan (Tokyo, Japan). Sense and antisense strands were hybridized and cloned into the pAAV-CMV-EmGFP-miRNA-backbone ^113^ digested with BsaI.

For construction of pCAG-S100a10 and pCAG-Htr1b, the S100a10 and Htr1b fragments were amplified by PCR from mouse cDNA obtained from DRN, and ligated to backbone obtained from pCAG-EGxxFP (addgene #50716).

For construction of pAAV-mTph2-EmGFP-miRNA-backbone, the mTph2 fragment from pAAV-mTph2-Venus-WPRE ^28^ and AAV backbone obtained from pAAV-CMV-EmGFP-miRNA-backbone ^113^ was ligated by using NEbuilder Assembly tool (https://nebuilder.neb.com). For construction of pAAV-mTph2-EmGFP-miRNA-Negative control, -*S100a10* and *-Htr1b*, the same Oligodeoxynucleotide primers as above were used. Sense and antisense strands were hybridized and cloned into the pAAV-mTph2-EmGFP-miRNA-backbone digested with BsaI.

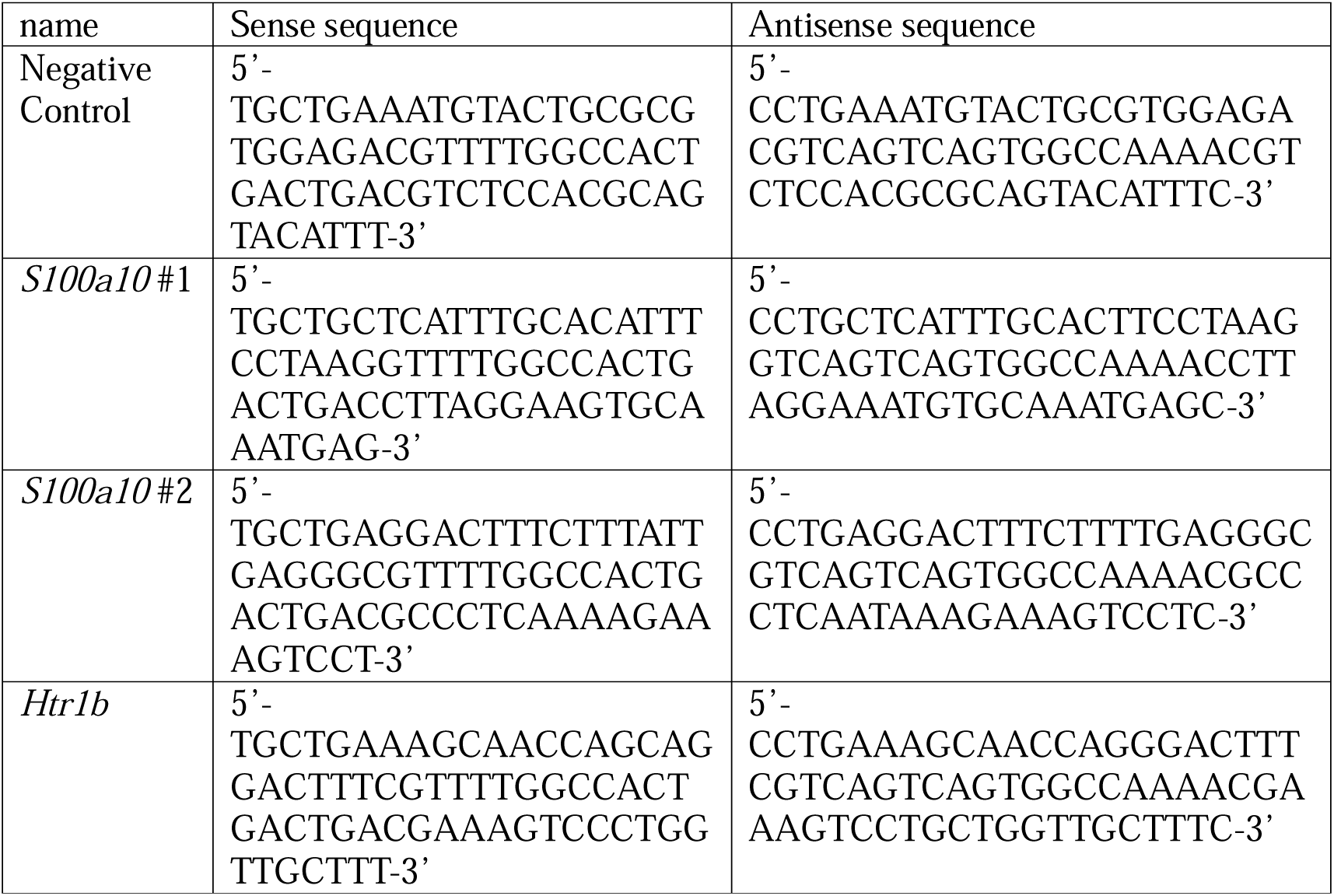

pAAV-mTph2-Venus-WPRE was the same as in the previous paper ^28,29^. For construction of pCAG-hSTAT6 (547V548T→AA, 641Y→F), wildtype STAT6 fragment was amplified by PCR from cDNA of HepG2 cells. Then the mutant STAT6 fragments were amplified by PCR from the wildtype fragment and ligated with CAG fragment from pCAG-EGxxFP (addgene #50716) by using NEbuilder Assembly tool. For construction of pAAV-mTph2-mSTAT6 (547V548T→AA, 641Y→F)-ALFA-WPRE, first wildtype STAT6 fragment was amplified by PCR from mouse cDNA obtained from spleen, and the ALFA-tag sequence ^114^ was inserted on the C-terminus. Then the mutant STAT6-ALFA fragments were amplified by PCR from wildtype fragment and ligated with mTph2 fragment from pAAV-mTph2-Venus-WPRE by using NEbuilder Assembly tool. For construction of pAAV-mTph2-SUN1sfGFP-WPRE, the SUN1sfGFP fragments were amplified by PCR from pAAV-Ef1a-DIO-Sun1GFP-WPRE-pA (addgene #160141), and ligated to backbone obtained from pAAV-mTph2-GCaMP6s-WPRE.

### Production and Purification of Adeno-Associated Virus (AAV) Vector

HEK 293T cells (Clontech, Mountain View, CA, USA) were grown in 15 cm dish to 60%–70% confluency, and 8 µg of pHelper, 5 µg of pAAV-DJ, and 5 µg of transfer plasmid were transfected with polyethylenimine (8mg/mL sterlile water, Polysciences, Warrington, PA, USA). After 60–72 h of incubation, the supernatant was aspirated and 500 µL of 1× Gradient Buffer was added to the cells on each plate, and then collected. The cell suspension was frozen in liquid nitrogen for 10 min, and placed in a 55 ◦C water bath until the cells were completely thawed. After lysis, they were triturated by using a 1 mL syringe and a 26-gauge needle (Terumo, Tokyo, Japan) for first two times, and vortexed for 2 min for last two times. This freeze-thaw cycle was repeated 4 times. After the addition of 0.5 µL of benzonase (Sigma-Aldrich, St. Louis, MO, USA), the lysate was incubated at 37 ◦C for 45 min, centrifuged for 15 min at 3000 × g in R20A2 rotor (Koki Holdings, Tokyo, Japan), and then the supernatant was collected. A discontinuous density gradient of 15%, 25%, 40%, and 58% iodixanol was prepared in an ultracentrifuge tube, and the supernatants were dripped onto the top layer of the density gradient. The tube was ultracentrifuged at 48,000 rpm, 18 ◦C for 1 h 45 min in a 50.2Ti rotor (Beckman-Coulter, Brea, CA, USA). After ultracentrifugation, a 1 mL syringe with a 26-gauge needle was inserted approximately 1–2 mm below the boundary surface between 40% and 58% gradient buffer layers, and 500 µL of solution was slowly extracted. This was aliquoted stored at −80 ◦C until use. Titlation of all AAVs were measured by qPCR (see Table below).

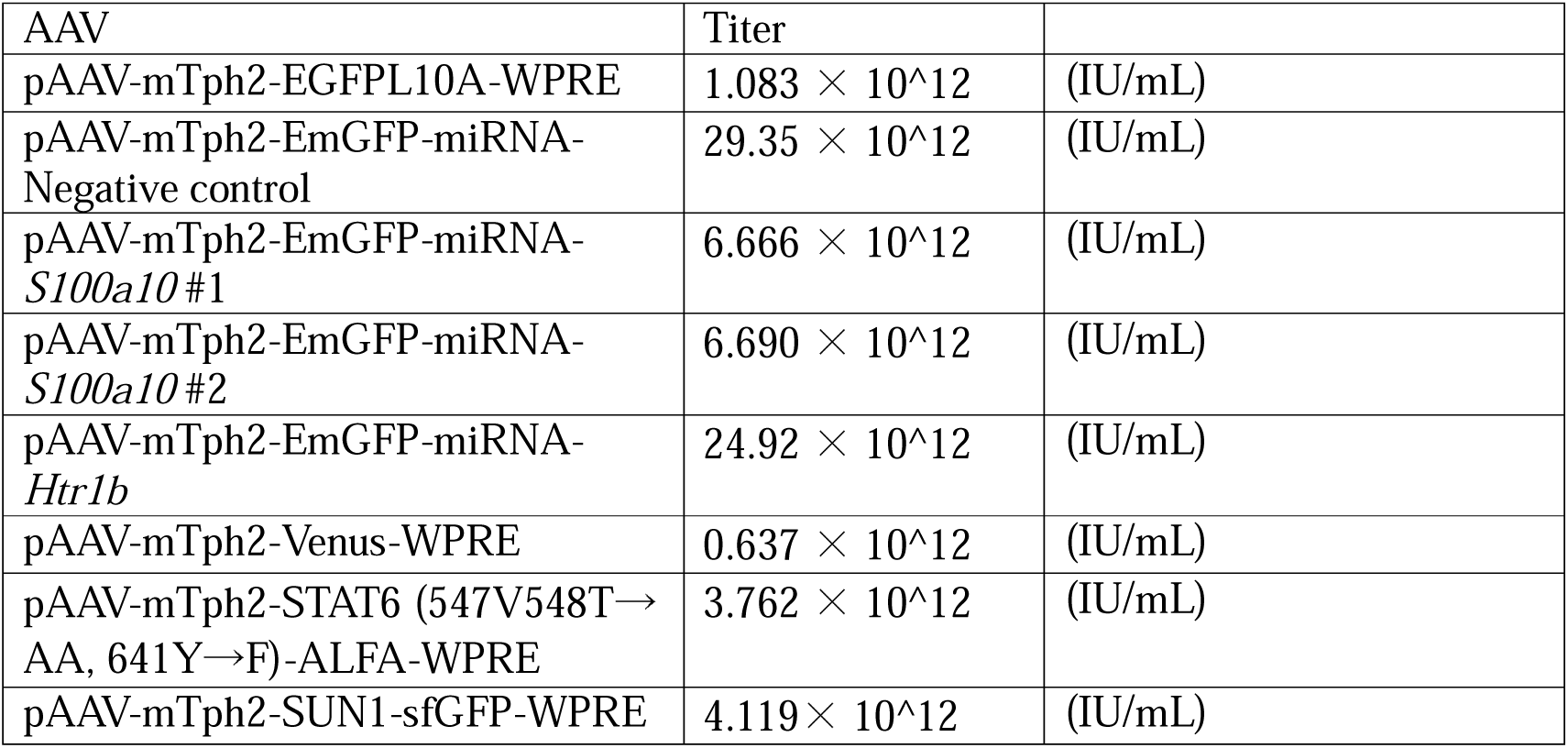

### Quantitative RT-PCR

The knockdown efficacy *in vitro* was measured by quantitative RT-PCR. First, HEK 293T cells (Clontech, Mountain View, CA, USA) were grown in 24 well dish to 50%– 60% confluency. Then 0.2 µg of a cloning plasmid (pCAG-S100a10 or pCAG-Htr1b), 0.6 µg of a knockdown plasmid (pAAV-CMV-EmGFP-miRNA-Negative control, pAAV-CMV-EmGFP-miRNA-*S100a10* #1, #2, or pAAV-CMV-EmGFP-miRNA-*Htr1b*) and 1.6 µL of lipofectamine 2000 (Thermofischer scientific, Carlsbad, CA, USA) were added into 100 µL of Dulbecco’s Modified Eagle Medium (DMEM). After 20 minutes incubation, the plasmid mixture was applied to the dish and incubated at 37 ◦C. After 6h the medium was changed and incubated again at 37 ◦C for 18-24h. After incubation, total RNA was isolated using ISOGEN reagent (Nippon Gene, Tokyo, Japan) in accordance with the manufacturer’s protocol, and cDNA was synthesized from 1□μg of total RNA using ReverTra Ace qPCR RT Master Mix (FSQ-201, Toyobo, Osaka, Japan). qPCR was performed using the StepOne real-time PCR system (Life Technologies, Carlsbad, CA, USA). The final reaction volume was 10□μl (5□ng of cDNA with primers and Thunderbird SYBR qPCR Mix, Toyobo, Osaka, Japan). The PCR conditions were as follows: heating for 1 min at 95 °C, followed by 35□cycles at 95□°C for 15□s and 60□°C for 1 min. The amount of human GAPDH in samples was used to normalize mRNA levels. Sequences of primers were following; hGAPDH Fw, ACTTCAACAGCGACACCCACT; hGAPDH Rv, GCCAAATTCGTTGTCATACCAG; S100a10 Fw, CACTGGTACCCCCACCCT; S100a10 Rv, TTGAGGGCAATGGGATGCAA; Htr1b Fw, GCCGACGGCTACATTTACCA; Htr1b Rv, GGTGATGAGCGCCAACAAAG.

### Translating Ribosome Affinity Purification (TRAP) and Microarray

TRAP for DRN serotonin neurons was performed as previously described ^24^ with minor modification. Briefly, mice were decapitated, and the DRN were rapidly dissected in ice-cold dissection buffer (1×HBSS, 2.5 mM HEPES, 35 mM glucose, 4 mM NaHCO3 and 0.1 mg/mL cycloheximide). DRN tissues from 2-3 mice were pooled for each experiment. Then DRN were transferred to ice-cold Lysis buffer (20 mM HEPES KOH, pH 7.3, 150 mM KCl, 10 mM MgCl2, 1% (vol/vol) Nonidet P-40 (NP-40; Nacalai Tesque, Kyoto, Japan), 0.5 mM DL-dithiothreitol (DTT; Sigma-Aldrich, St. Louis, MO, USA), 0.1 mg/mL cycloheximide, EDTA-free protease inhibitors (Roche, Basel, Switzerland) and 10 μL/ml RNasin® Plus Ribonuclease Inhibitor (Promega, Madison, WI, USA). The tissue lysates were then homogenized on ice and 1% of each lysate volume was separated as Input samples. Post-nuclear supernatant was prepared by centrifugation at 2,000 g for 10 min at 4 ◦C. The S20 fraction was prepared by adding 300 mM 1,2-diheptanoyl-sn-glycero-3-phosphocholine (DHPC; Avanti Polar Lipids) to the supernatant followed by incubation on ice for 5 min and centrifuged at 20,000 g for 10 min at 4 ◦C. Immunoprecipitation of ribosomes was carried out by incubation of the supernatant with DynabeadsTM Protein G (Invitrogen) and anti-GFP monoclonal antibodies (19C8 and 19F7; Memorial Sloan Kettering antibody facility) at 4 ◦C with end-over-end rotation for 16 h. Beads were collected on a magnetic rack, washed with high-salt buffer (20 mM HEPES-KOH, pH 7.3, 350 mM KCl, 10 mM MgCl2, 1% NP-40, 0.5 mM DTT, and 100 mg/ml CHX) and immediately subjected to RNA extraction using absolutely RNA nanoprep kit (Agilent Technologies) with in-column DNase digestion. The quality of purified RNAs was evaluated based on RNA integrity numbers (RIN, Schroeder et al., 2006) read by an Agilent Technologies BioAnalyzer 2100. Complementary DNA (cDNA) libraries were prepared according to the GeneChip WT Pico Kit (Thermofischer scientific). Then Clariom S Assay was performed according to the manufacturer’s instruction (Affymetrix).

### Gene expression analysis

Microarray data were analyzed using the Affymetrix Transcriptome Analysis Console 4.0 (TAC 4.0, Affymetrix) to identify differentially expressed genes. For calculate fold change differences between samples, duplicate genes were filtered out in advance using Python 3 and 21,905 genes were used for the subsequent analysis. For heatmap visualization, log-transformed gene expression was further standardized (average = 0, variance = 1), and the standardized values of Z scores across samples were visualized as a heatmap using Prism 9. Principal component analysis (PCA) was conducted using R package DESeq2 ^116^.

### IPA analysis

List of genes that were either significantly upregulated or downregulated in SSRI treated mice and CSDS susceptible mice compared with each control (that is, naïve mice and CSDS resilient mice respectively) were analyzed using the Ingenuity Pathway Analysis (IPA) canonical pathway and upstream regulator analysis (Qiagen Ingenuity Systems, Venlo, Netherland). Gene expression changes with absolute values greater than 2 were considered significant.

### Stereotaxic surgeries

Stereotaxic surgeries were conducted using a small animal stereotaxic frame (Narishige, Tokyo, Japan) according to the Brain Atlas (Franklin & Paxinos, 2007). The animals were anesthetized with a combination of 0.3 mg/kg of medetomidine, 4.0 mg/kg of midazolam, and 5.0 mg/kg of butorphanol. The adeno-associated virus (AAV) vectors were microinjected at 1 μL per mouse into the DRN (AP –4.4 mm, ML +1.1 mm, depth +3.7 mm at 20◦ from the bregma,). We used AAV-DJ encoding Venus (mTph2-Venus-WPRE), a GFP-ribosomal fusion protein EGFPL10A (mTph2-EGFPL10A-WPRE), a miRNA control fused with EmGFP (mTph2-EmGFP-miRNA Negative Control), a miRNA for *S100a10* fused with EmGFP (mTph2-EmGFP-S100a10 miRNA RNAi #1/#2), a miRNA for *Htr1b* fused with EmGFP (mTph2-EmGFP-Htr1b miRNA RNAi), or a mutant of STAT6 (mTph2-STAT6 (547VT548→AA, 641Y→F)-ALFA-WPRE), a SUN1 fused with sfGFP (mTph2-SUN1sfGFP-WPRE) under the control of the mouse Tph2 promotor. 2 or 4 weeks after viral injection, the animals were used for the knockdown experiments or mutant experiments respectively. After behavioral analyses, all mice were sacrificed, and the AAV infection was verified immunohistochemically. For IL4 microinjection, Animal-Free Recombinant Murine IL-4 (AF-214-14, Peprotech, Cranbury, NJ, USA) was diluted to 10 pg/μL or 100 pg/μL with saline and microinjected at 1 μL per mouse into the DRN same as the viral injection. Some animals were sacrificed 45 min after injection to confirm whether IL4 microinjection increased pSTAT6, and the rest was used for the following behavioral analysis.

### Chronic Social Defeat Stress

4 weeks after stereotaxic surgery, chronic social defeat stress was applied as described previously ^32^ with minor modifications. Briefly, ICR mice were screened based on their aggressiveness to a naïve C57BL/6J mouse, as measured by the latency and the number of attacks, and were used as aggressor mice for chronic social defeat stress. Before the beginning of chronic social defeat, mice were isolated for 1 week (in other words, isolation was conducted 3 weeks after the stereotaxic surgery). Food and water were available *ad libitum*. For chronic social defeat, an isolated mouse to be defeated was introduced and kept in the homecage of a resident aggressor ICR mouse for 5 min daily for 10 consecutive days. The pair of defeated and aggressor mice was changed daily to minimize the variability in the aggressiveness of ICR. The social interaction test was performed at the next day of final defeat session as described below. The social interaction test was performed as described previously ^32^ with minor modifications. Individually housed mice were acclimated to testing rooms under the red dim light for at least 30 min before the test. First, for habituation to a test environment, a defeated mouse was kept for 150 sec in an open field chamber (50 × 50 × 50 cm) with an empty wire mesh cage (10 × 6.5 cm) located at one end of the field. Consecutively, the same mouse was kept for 150 sec in the same open field chamber with an unfamiliar ICR mouse enclosed in the wire mesh cage. Mouse behaviors were video monitored, and the trajectory of mouse ambulation was determined and recorded by ANY-MAZE (Stoelting Co., Wood Dale, IL, USA). The four corners of the chamber (9 cm square) defined as the avoidance zone and the area surrounding the wire mesh cage (14 × 24 cm) defined as the interaction zone. Interaction ratio was calculated as time spent in the interaction zone in the presence of an unfamiliar ICR mouse divided by that in the absence of ICR mouse. The proportion of time spent in the interaction and avoidance zones during the observation period (150 sec) and interaction ratio was used as an index for the level of social avoidance.

### Behavioral Analysis

Behavioral tests for evaluation of an antidepressant-like effect SSRI were conducted from the day after the SSRI administration was ended. We used both group-housed mice and single-housed mice in this experiment. Tail suspension test was conducted at first and the open field test was conducted with at least 24 hr of interval.

For viral injection experiments and IL4 microinjection experiments, behavioral tests were conducted two weeks after stereotaxic surgery except for the experiment of STAT6 mutation. The analysis of STAT6 mutation was conducted four weeks after stereotaxic surgery. Multiple tests were conducted in the following order: Open field test, elevated plus maze test, and tail suspension test, with at least 24 hr of interval between tests.

#### Open Field Test

Locomotor activity was measured using an open field test. The open field arena consisting of a white acrylic cube (50 × 50 × 50cm) was used. Each mouse was placed in the center of the open field apparatus and permitted free exploration. The behavior of mice was recorded with a camera over a 10-min session. Total distance traveled were analyzed automatically using video tracking system (ANY-maze version 4.99 or 6.0, Stoelting, Wood Dale, IL, USA).

#### Elevated Plus Maze Test

The apparatus consisted of two open arms and two closed arms (30 × 5 cm) extended from a central platform (5 × 5 cm). Each animal was placed individually into the central platform. The behavior of the animal was recorded with a camera over a 10-min session. Time spent in the open and closed arms and total distance traveled during a session were measured automatically using video tracking system (ANY-maze version 4.99 or 6.0). Percentage of the time spent in the open arm was calculated as following; 100 × open arm time (s) / (open arm time (s) + closed arm time (s)). The latency to first entered either arm was calculated as the time spent in the open arm.

#### Tail suspension Test

The tail suspension test was performed as previously described ^118^ with brief modifications. Briefly, distal end of the tail was taped to a force-transducer (PowerLab 2/26, AD Instruments, Colorado Springs, CO, USA) fixed to the ceiling of a test box (40 × 40 × 40 cm). Mice were suspended for 6 min, and total immobile time was calculated from the trace of applied force. The behavior of each mouse was manually evaluated during test session by the experimenters, and mice that held their hindlimbs or climbed their tails with their forelimbs were excluded from the analysis.

### Histology

The animals were deeply anesthetized with a combination of 0.3 mg/kg of medetomidine, 4.0 mg/kg of midazolam, and 5.0 mg/kg of butorphanol and transcardially perfused with PBS followed by 4% paraformaldehyde (Nacalai Tesque, Kyoto, Japan) in 0.1 M PBS. After perfusion fixation, the brains were harvested, equilibrated in 15% sucrose in PBS overnight and frozen at −80[◦C. The brains were cryosectioned into 30 µm-thick coronal sections with the cryostat (Leica CM3050S; Leica Biosystems, Nussloch, Germany) and stored at −80 ◦C until immunohistochemical processing. For immunohistochemistry, the DRN sections were first washed three times with PBS containing 0.1% Triton X-100 for 5 min each time and treated with the blocking buffer (PBS containing 0.25% Triton X-100 and 5% horse serum) for 1 h at room temperature for permeabilization and blocking. Then sections were incubated overnight at 4 ◦C with goat polyclonal anti-S100a10 antibody (1:200; AF2377, R&D Systems, Minneapolis, MN, USA), rabbit polyclonal anti-5HT_1B_ Receptor antibody (1:100; ab13896, abcam, Cambridge, UK), sheep polyclonal anti-tryptophan hydroxylase (TPH) antibody (1:500; AB1541, Merck Millipore, Burlington, MA, USA), and rabbit polyclonal Anti-STAT6 (phospho Y641) antibody (1:100; ab28829, abcam) diluted in the blocking buffer. GFP fluorescence was observed without using antibody. After incubation, sections were washed in PBS three times and incubated with Alexa Fluor 594-labeled donkey anti-goat IgG (1:200; A11058, Life Technologies, Carlsbad, CA, USA), Alexa Fluor 594-labeled donkey anti-rabbit IgG (1:200; A21207, Life Technologies), or Alexa Fluor 594-labeled donkey anti-sheep IgG (1:200; A11016, Life Technologies) for 2h at room temperature. Then the sections were washed in PBS and mounted on glass with DAPI Fluoromount-G (Southern Biotechnology, Birmingham, AL, USA). Coimmunostaining of pSTAT6 and Tph2 performed the same as above, except that the concentration of Triton X-100 in blocking buffer was set to 1%. For coimmunostaining of S100a10 and Tph2, mouse monoclonal anti-tryptophan hydroxylase (TPH) antibody (1:1000; AMAB91108, Sigma-Aldrich, St. Louis, MO, USA) was used instead of the anti-sheep antibody. Immunofluorescence was visualized using a laser scanning confocal microscopy (Fluoview FV10i, Olympus, Tokyo, Japan) with software (FV10i-SW, Olympus, Tokyo, Japan; Image J, NIH, Bethesda, MD, USA).

### Image analysis

For counting the number of Tph2, GFP, S100a10 positive cells (corresponding to Extended Data Figure 1a, 4, 7), the average number of two images per animals were used. For calculating S100a10 or 5-HT_1B_R intensity (corresponding Figure 2d, e, h, 3a, d, 5c), images were analyzed using ImageJ 1.53a (National Institutes of Health, USA, http://imagej.nih.gov/ij). Intensity of each background (an area without any positive signal, 32 × 32 pixels) was subtracted from the intensity of each full images (1024 × 1024 pixels). Also, the number of DAPI for these corresponding images was calculated. For quantitative analysis of pSTAT6 (corresponding to figure 4d), Tph2-positive and DAPI-positive regions were extracted using ImageJ and the number of dots in those regions were counted.

### Western Blotting

Immunoblotting analyses were conducted on whole-cell lysates, as previously described with minor modifications ^119^. Protein samples were quantified by Pierce BCA protein assay kit (Thermofischer Scientific) and 20 μg of total protein were loaded onto a 14.5% (for S100a10) or 10% (for pSTAT6, STAT6) SDS-polyacrylamide gel and blotted onto ClearTrans® PVDF Membrane (FUJIFILM Wako Pure Chemical Corporation, Osaka, Japan). After blocking with Blocking One (for S100a10, STAT6, GAPDH; Nacalai Tesque, Kyoto, Japan) or Blocking One P (for pSTAT6; Nacalai Tesque, Kyoto, Japan), membranes were incubated overnight at 4 °C with the following primary antibodies: rabbit anti-pSTAT6 (1:1000, ab28829, abcam), rabbit anti-STAT6 (1:1000, #5397, Cell Signaling Thechnology), mouse anti-GAPDH (1:1000, sc-32233, Santa Cruz Biotechnology, Dallas, TX, USA), or goat anti-S100a10 (1:200, AF2377, R&D Systems). On the following day, the membranes were incubated 1h at room temperature with following horseradish peroxidase (HRP)-conjugated secondary antibody: HRP-conjugated anti-rabbit IgG (1:5000, NA934V, Cytiva, Tokyo, Japan) or HRP-conjugated anti-mouse IgG (1:10000, 115-035-003, Jackson Immunoresearch, West Grove, PA, USA). Note that STAT6 was immunoblotted after stripping the pSTAT6 signal by using WB Stripping Solution Strong (Nacalai Tesque, Kyoto, Japan) according to the manifacture’s guide. The blots were visualized using an ECL system (Tanon 4600, Shanghai, China) and analyzed by the ImageJ software.

### ELISA

Brain sampling for ELISA was conducted according to the previous report ^120^ with some minor modification. Briefly, brains were rapidly removed after cervical dislocation and washed gently with ice-cold PBS and sliced to 1-mm thick coronal sections using the mouse brain slicer matrix on ice. Then the DRN was dissected using a razor. Samples were collected into 1.8 mL Nunc cryotubes (Thermo Scientific), immediately frozen in liquid nitrogen, and stored in liquid nitrogen tank until use. To measure brain IL-4 and IL-13 level, 150 µL extraction solution (20 mmol/L Tris-HCl, 150 mmol/L NaCl, 1 % TritonX-100, 1 mM EDTA and 1×Complete EDTA free-tablet (Roche Diagnostics, Basel, Switzerland) in distilled water) was added each frozen brain sample, and samples homogenized on ice. Samples were nutated for 90 min at 4 °C, then centrifuged at 1000 × g for 20 min at 4 °C, and the supernatants were collected into new tube. They were stored at −80 °C until ELISA assays were performed.

Frozen samples were thawed and brought to room temperature. Cytokine levels were measured using Mouse IL-4 Quantikine ELISA (M4000B; R&D Systems) and Mouse IL-13 Quantikine ELISA (M1300CB; R&D Systems). 50 μL of samples were used for each measurement and samples were measured as single samples due to the small volume. Plates were read on microplate reader (model 680; Biolad, Hercules, CA, USA). Total protein concentration was measured using Pierce BCA protein assay kit and cytokine concentration was expressed as a % of total protein.

### Luciferase assay

HepG2 cells were grown in Dulbecco’s modified Eagle’s medium (Nacalai Tesque) supplemented with 10% heat-inactivated fetal bovine serum (SigmaAldrich), 100 U/mL penicillin (Nacalai Tesque), and 100 μg/mL streptomycin (Nacalai Tesque). 475 ng of the reporter plasmid (p4xSTAT6-Luc2P (addgene #35554)) and 475 ng either of pCAG-hSTAT6 (VT→AA, Y→F) or pcDNA was transfected by using PEI MAX (Polysciences, Warrington, PA, USA) according to the manifacture’s guide. 42 h following transfection, the cells were stimulated with 10 ng/ml IL-4. Then 48 h following transfection (namely, 6 h after IL-4 stimulation), 50 μL of PicaGene Luminescence Kit (TOYO B-NET, Tokyo, Japan) was added for each well and the luciferase activity was immediately observed by Spark^®^ multimode microplate reader (TECAN, Männedorf, Switzerland).

### Statistical analysis

Statistical analysis was conducted using Graphpad Prism 9 (GraphPad, San Diego, CA, USA). Briefly, one-way ANOVA with Dunnett’s *post hoc* test was used for comparisons between multiple experimental groups. Unpaired *t-*test or unpaired t-test with Welch’s correction were used for comparison between two groups according to the results of the F test. Values in text and graphs were expressed as mean ± SEM. All *P*-values reported are two tailed. Statistical significance was defined as *P* < 0.05. The numbers of animals used in each experiment are indicated in the figure legends.

